# Smartphone Placement Recognition during Walking: Performance Determinants and Real-World Generalizability

**DOI:** 10.64898/2026.05.12.724503

**Authors:** P. Tasca, G. Trentadue, E. Buckley, S. Sun, M. Long, N. Ireson, F. Ciravegna, V. Lanfranchi, A. Cereatti

## Abstract

The opportunity to collect movement data from smartphones for prolonged periods has opened new perspectives in the field of clinical movement analysis. However, when monitoring people’s mobility in free-living conditions, smartphone placement can influence the validity of the extracted digital mobility outcome. This study aimed to develop and validate an automatic smartphone placement recognition classifier and to investigate potential critical factors that can influence performance.

The classifier was trained on data from 15 healthy participants using inertial signals collected from smartphones placed at six body placements during free-living walking and externally validated on over 3,000 individuals from external datasets, including blind participants and patients with cardiovascular or Parkinson’s disease. A decision-tree ensemble model was developed using feature subsets of increasing dimensionality, with the optimal subset comprising 50 features.

Classification accuracy increased consistently when front and back pocket placements were aggregated (81.1%) and further improved when coat pocket was also included in the pocket class (88.5%), underscoring the challenge of distinguishing between fine-grained pocket placements. The best-recognized placements across the external datasets were lower back (precision: 100%, recall: 72.5%), hand (precision: 94.2%, recall: 94.5%), and the aggregated pocket class (precision: 86.7%, recall: 90.2%). Recognition accuracy changed across cohorts (0.73 – 0.85), activities (0.63 – 0.94) and speed (0.79 – 0.87), however it stayed consistent across various technological and environmental factors.

Overall, this study demonstrates the feasibility of robust placement recognition in walking and underscores the importance of accounting for key influencing factors when designing frameworks intended for deployment in heterogeneous real-world or clinical contexts.

**Highlights:**

- Machine learning accurately identifies smartphone placement during real-world gait
- Six on-body placements recognized, including pockets, hand, bag, and lower-back
- Free-living data used for training, ensuring robust performance across conditions
- Feature selection and hyperparameter tuning optimize classification accuracy
- External validation confirms generalizability across >3,000 healthy and diseased adults

## 1. Introduction

Smartphones, with their embedded inertial measurement units (IMU), are used daily by people, making them the ideal tool for assessing digital mobility and gait [1]. Most IMU-based algorithms for mobility assessment in research and clinical applications assume a fixed device placement, typically the lower back [2], the hand [3] or pockets [4]. However, in real-world settings, on-body placement varies continuously based on user habits, gender, and age [5], compromising the reliability of spatio-temporal mobility metrics [6]. Automatic smartphone placement recognition is therefore a required preliminary step to select the most appropriate algorithms and to improve clinically and ecologically valid digital mobility assessment [7].

Several IMU-based smartphone placement recognition methods have been proposed including decision trees [8], support vector machines [9], heuristic methods [10], ensemble models [11] and deep-learning [12], reporting accuracies between 85% and 99%. However, only few studies evaluated smartphone placement recognition classifiers on a small number of external datasets [7], [13], [14], and none systematically examined how potentially critical contextual and subject-related factors influence generalization, limiting understanding of how smartphone placement recognition algorithms perform in real-world scenarios.

The aims of this study were: (1) to develop a smartphone placement recognition classifier to recognize six on-body smartphone placements during real-world walking; (2) to validate it on unseen data from over 3,000 healthy and diseased individuals, including blind people, cardiovascular, and neurological patients; and (3) to assess how contextual and subject-related factors affect performance.

The final goal was to evaluate the robustness of smartphone placement recognition methods using heterogeneous datasets, providing a systematic assessment of their generalizability across diverse conditions.

## 2. Materials and Methods

### 2.1 Analysed Datasets

This study utilised both new data collected during this study and fifteen external datasets available online.

#### 2.1.1 New Datasets

Smartphone placement recognition was framed as a multi-class classification problem involving six common smartphone placements [5]: Hand-held, Shoulder Bag, Front Pocket, Back Pocket, Coat Pocket, and Lower-Back (Figure 1A).

**Figure 1.**
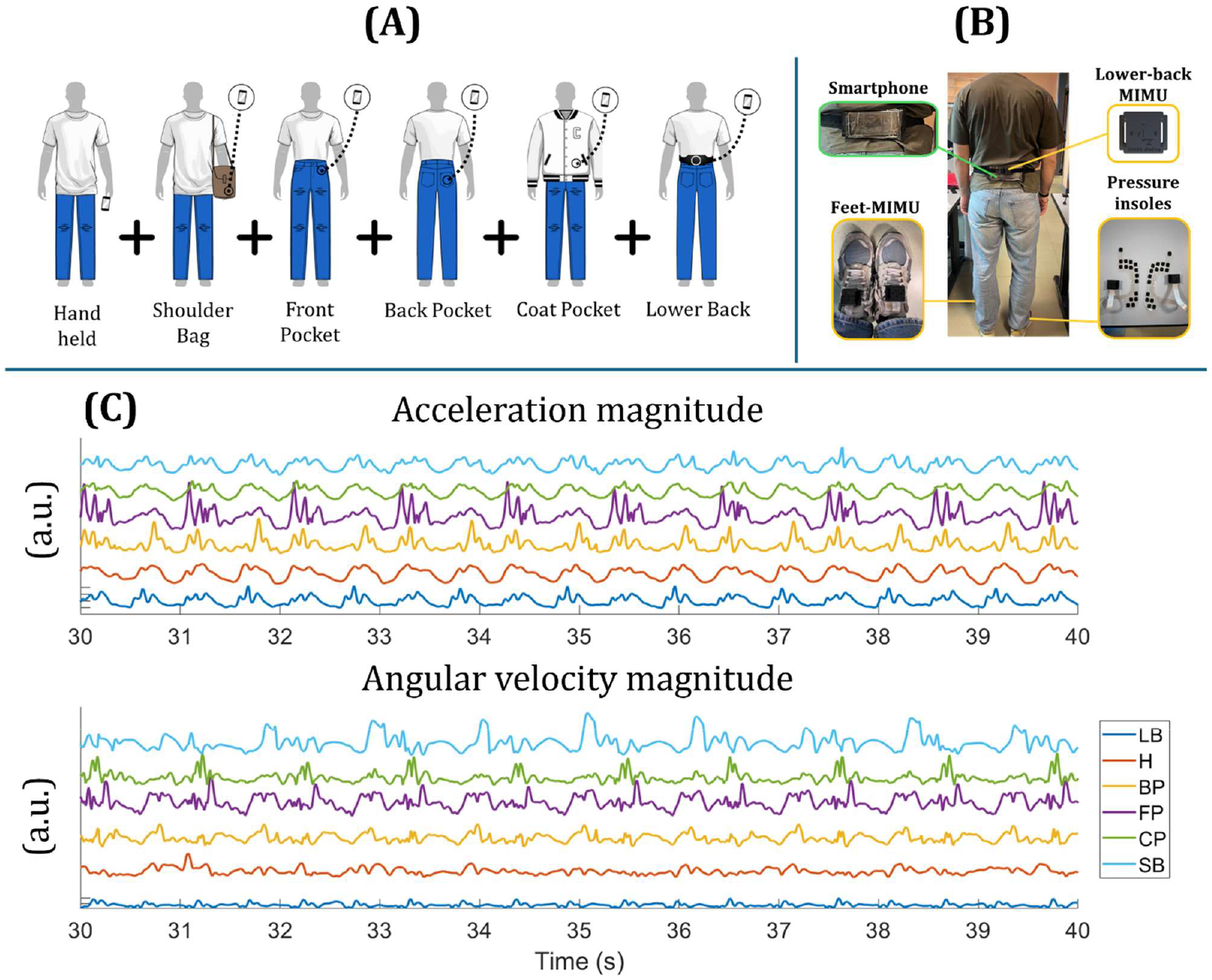
A) Schematic representation of smartphone placements included in the recognition framework. During acquisitions, all smartphone placements recorded data simultaneously. Hand-held (H): Held in the hand during the natural arm swing motion; Shoulder Bag (SB): Carried in a shoulder bag, Front Pocket (FP): In the front pocket of the trousers or jeans; Back Pocket (BP): In the back pocket of the trousers or jeans; Coat Pocket (CP): In the side pocket of a jacket or coat; Lower-Back (LB): Belt-fixed at the level of the 5th lumbar vertebra with landscape orientation. B) Overview of a participant equipped with the multi-sensor reference system, including magneto-IMUs on the feet and the lower-back at L5 and piezoresistive pressure insoles. The LB smartphone and the lower-back MIMU of the reference system were co-located at the L5 vertebra to facilitate synchronization between the two systems. C) Representative examples of accelerometer and gyroscope magnitude signals recorded from smartphones across the six different placements during a 10-second walking window. Signals are offset vertically for clarity. a.u.: arbitrary units.

Inertial data were collected at 100 Hz with six Samsung Galaxy A34 smartphones through a custom application. The zero-level accelerometers (-0.04 ± 0.02 m/s^2^) and gyroscopes (0.01 ± 0.06 deg/s) errors were quantified through static calibration [15]. Analysis was restricted to walking segments, identified by the INDIP multi-sensor system [16], [17]. Smartphone and INDIP timestamps were aligned through cross-correlation analysis (Figure 1B).

Data was collected in laboratory and free-living conditions from 15 healthy participants (10 males, 5 females, 22-57 y.o):

- **New Laboratory Dataset:** straight-line and oval walking paths in a corridor at three self-selected speeds (slow, regular, fast), each repeated thrice.
- **New Free-living Dataset:** A 2.5-hour free-living acquisition in the city of Sheffield (UK).

Participants were equipped with the INDIP system and the six smartphones in the defined placements with unconstrained orientation, to preserve real-world orientation variability. The protocol was approved by the University of Sheffield Ethics Committee (application no. 064746).

#### 2.1.2 Existing External Datasets

External validation was conducted using fifteen existing datasets comprising smartphone-recorded walking data acquired at known body placements [5]. Only datasets that were publicly available or accessible upon request and included both acceleration and angular velocity signals were considered (Figure 2). In total, approximately 250 hours of walking data from more than 3000 participants were analyzed. Smartphone placements not matching those of the New Datasets were excluded.

**Figure 2.**
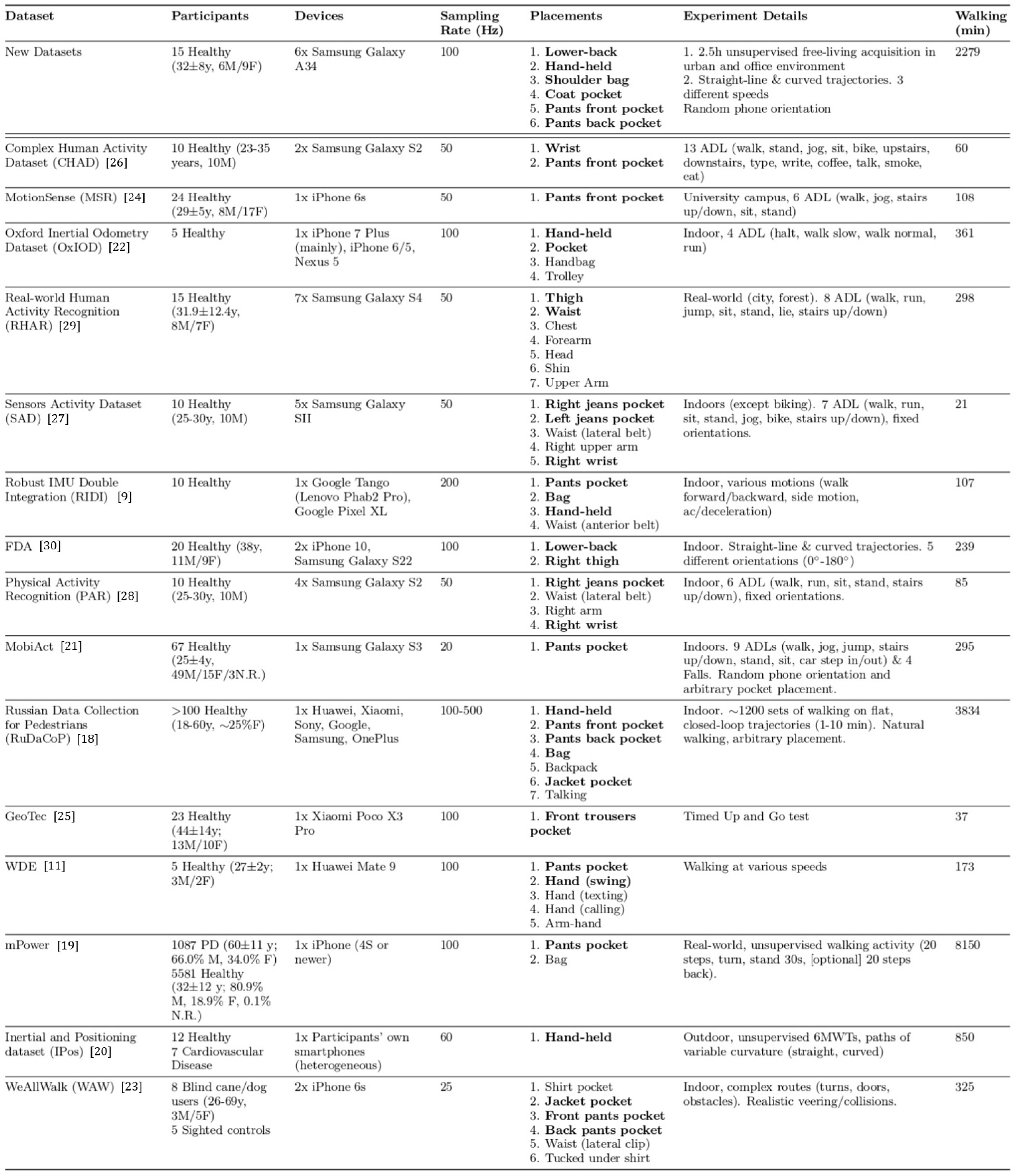
Summary of the datasets used for validation in the present study. PD: Parkinson’s disease; ADL: Activities of Daily Living; 6MWT: 6 Minutes Walking Test; N.R.: Not Reported. i. Smartphone placements matching the ones included in the framework are highlighted in bold. ii. The number of smartphones used for a single recording is reported in the “Devices” column. iii. The column “Walking (min)” shows the total duration of walking data recorded by all the equipped devices. iv. For IPos, IMU data was available for 15 participants out of 19, the dataset version used corresponds to the November 2024 release. v. For WAW, 10 blind participants instead of 8 were retrieved from the online data repository. Of them, 3 walked using a guide dog, and 9 using a walking cane. vi. For mPower, only 2245 healthy and 665 PD patients were retrieved from the public data repository based on the reported “Professional Diagnosis” variable. Most PD patients were tested before and after medication. No information about disease subtypes was available in the dataset. The dataset version used corresponds to the latest update available as of January 2025.

### 2.2 Development and Validation of the smartphone placement recognition framework

All the processing steps were performed using MATLAB R2024b (The Mathworks, Inc.).

#### 2.2.1 Feature Extraction

All data was resampled to 100 Hz. Walking intervals were extracted either using the INDIP system for the New Datasets or annotated labels for External Datasets. Acceleration and angular velocity magnitudes [31] were segmented into 10-s windows, with shorter recordings circularly padded to preserve signal periodicity. Stationary windows (mean angular velocity < 20 deg/s) were removed. To account for placement-specific patterns (Figure 1C), over 150 features from time, frequency, energy, entropy, and wavelet domains were extracted.

#### 2.2.2 Smartphone Placement Recognition Classifier

A multi-class tree-ensemble classifier was trained using MATLAB’s *fitcensemble* function with AdaBoost [32], selected for its strong performance in prior smartphone placement recognition studies [29], [33]. A two-stage feature selection procedure was applied: first, features were ranked according to the Minimum Redundancy Maximum Relevance criterion [34], second, a wrapper-based approach was applied, iteratively training models using progressively larger feature subsets, ranging from 10 to 156 features in increments of 10.

The New Free-living Dataset (15 healthy participants) was randomly split into a Training Set (10 participants) and a Test Set (5 participants). For each feature subset size, Bayesian optimization (50 iterations) was applied on the Training Set to tune the ensemble hyperparameters and identify the best-performing model. The best-performing models obtained for each feature subset were compared on the Test Set to identify the optimal feature subset size. The model trained using the optimal subset was subsequently adopted as the final model for all downstream validation analyses.

The code and the extracted features, together with detailed information on feature selection, hyperparameter optimization, and model training procedures, are publicly available on GitHub at: https://github.com/H-MOVE-LAB/smartphone-placement-recognition

#### 2.2.3 Evaluation Framework

The classifier was trained using six placement classes to reflect the diversity of smartphone placements and to explicitly assess discrimination between biomechanically-similar pocket-related configurations, as well as more challenging configurations such as shoulder bag. Training on the full set of classes ensured that placement-specific inertial patterns were learned without simplifications during model construction.

To quantify the impact of misclassification among pocket-related placements, additional classification settings were analysed at the validation stage only, by aggregating pocket classes while keeping the trained model unchanged. Specifically, performance was assessed on three classification settings (Figure 3):

**Figure 3.**
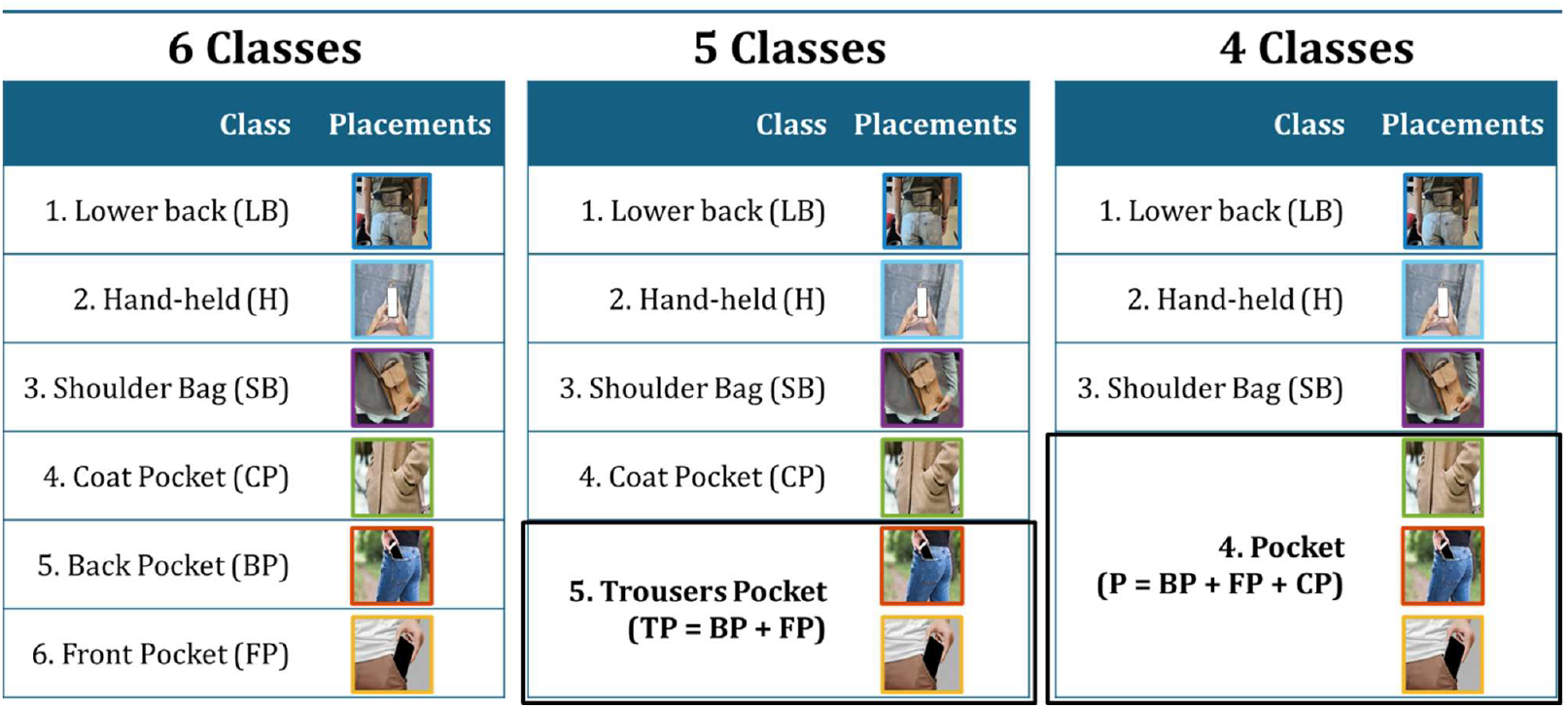
Schematic representation of the three smartphone placement classification settings, illustrating the original 6-class configuration and the progressive aggregation of pocket-related placements into 5-class and 4-class classification schemes.

- 6-class setting: all six smartphone placements treated as separate classes.
- 5-class setting: Front and Back Pocket merged into a single class.
- 4-class setting: Front, Back, and Coat Pocket merged into a single class.

This strategy enabled assessment of performance changes when progressively relaxing distinctions between pocket-related placements, without retraining the model, and facilitated comparison across datasets with heterogeneous or unspecified pocket positioning.

Performance evaluation of the final model included internal and external validation on fully unseen data (Section 2.1.2). Internal validation was performed on the Test Set (data from 5 participants of the New Free-living Dataset not used for training) and the New Laboratory Dataset. External validation was carried out on the 15 External Datasets (Figure 2). Both validation procedures were performed consistently across all three classification settings, obtained by aggregating pocket-related classes as described above.

Performance metrics included overall accuracy, per-class recall, and per-class precision, defined from true positives (TP), false positives (FP), and false negatives (FN) for each class.

Performance was assessed across the different classification settings and, across increasing feature subset dimensionality, using exclusively unseen data.

To further evaluate robustness, a dedicated analysis examined the influence of demographic, hardware, and contextual factors on classification performance. This analysis focused on overall recognition accuracy averaged among the three most represented placements in the External Datasets - Hand, Lower-Back, and Pocket (aggregating Back, Front and Coat Pocket) - while excluding less frequent and more challenging placements (e.g., Shoulder Bag), which were analysed separately to avoid masking factor-specific effects. Analysed factors included activity, morbidity, gender, age, hardware, sampling frequency, walking trajectory, and walking speed. For each factor, appropriate clusters of datasets were selected. Each factor was investigated in isolation by controlling all others, using appropriate dataset clusters. Further details are available in the accompanying GitHub repository.

## 3. Results

### Feature Selection results

Feature selection analysis showed a consistent increase in accuracy across all three classification settings as feature dimensionality increased up to the top-50 ranked features, after which performance reached a plateau (77.3%, 81.1%, and 88.5% accuracy for the 6-, 5-, and 4-class settings, respectively), suggesting that including additional features did not yield meaningful improvements (Figure 4). Hyperparameter optimization with this feature subset yielded a tree ensemble with 449 learners, 362 maximum splits, and 0.82 learning rate.

**Figure 4.**
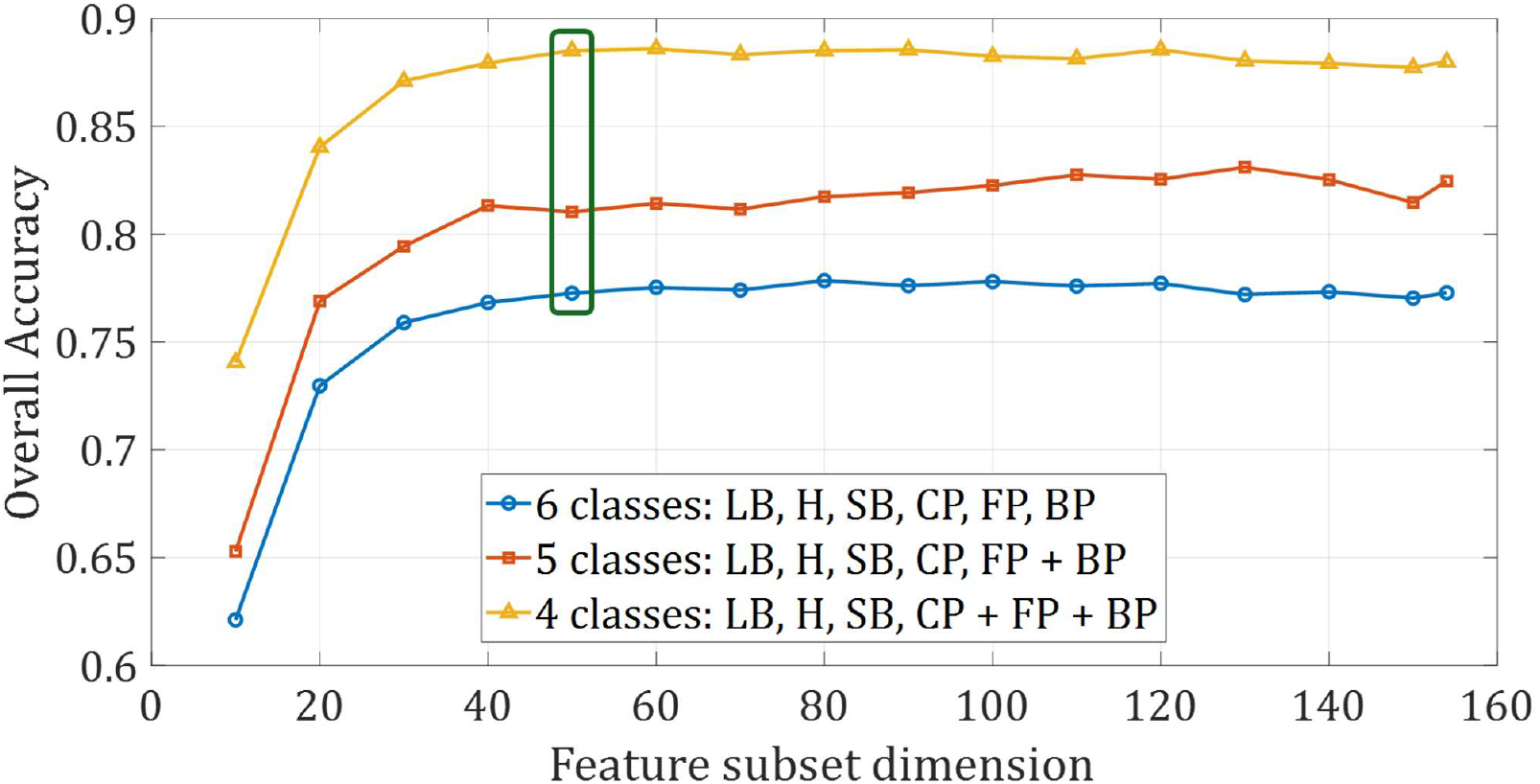
Comparison of overall accuracy achieved by the ensemble classifier among the 6-class, 5-class, and 4-class settings across increasing number of features (feature subset dimension). Performance plateaued at the top 50-ranked features, yielding accuracies of 77.3%, 81.1%, and 88.5% in the 6-, 5-, and 4-class settings, respectively. Accuracy is calculated on data of the 5 participants of the New Free-living Dataset not used for training the model. The optimal feature subset was determined by selecting the feature subset size for the 4-class setting where the accuracy increment over the preceding, smaller subset was at least 0.5%.

### Internal Validation Results

Internal validation of the optimal model on the Free-living Test Set provided an estimate of classifier performance on data homogeneous with training data, thus representing a reference performance under near-optimal conditions. Hand and Shoulder Bag placements achieved the highest recognition performance, with precision ranging between 88–96% and recall between 87–95% (Figure 5). In contrast, front pocket was the poorest-recognized class, exhibiting a markedly low recall (26.6%). This limitation was substantially mitigated by aggregating pocket-related classes: merging back and front pockets increased recall to 61.4% with a precision of 93.3%, while further aggregating coat pocket yielded a recall of 84.0% and a precision of 94.7%. The model also demonstrated good generalizability when evaluated on data acquired under controlled conditions (New Laboratory dataset), achieving consistent performance across all three classification settings (precision: 79–84%; recall: 79–87%). Additional insights into the structure of misclassification in the Free-living Test Set are provided in the Supplementary Materials.

**Figure 5.**
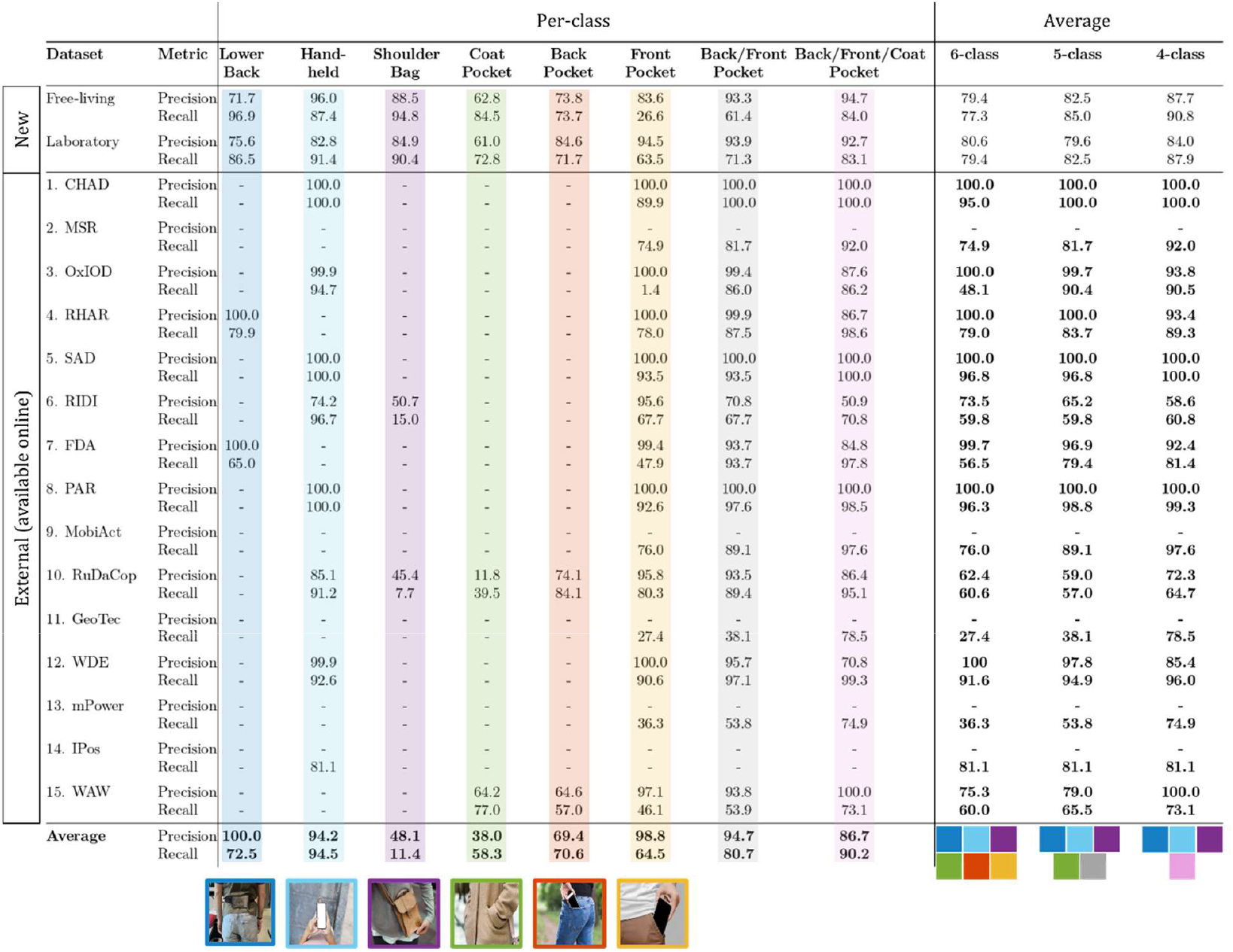
Per-class precision and recall are reported for each dataset and smartphone placement, including the six original classes and the aggregated pocket classes associated with the 5-class (Back/Front Pocket) and 4-class (Back/Front/Coat Pocket) classification settings. Columns are color-coded to identify each placement, with the corresponding class symbol reported below each column. For each dataset, average performance values are shown together with colored markers indicating the number and type of smartphone placements included in the averaging process, depending on the classification setting. For datasets including only one placement, per-class precision would trivially equal 100% and was therefore not reported. Average recall (reported in the “Average” columns) corresponds to balanced accuracy, computed as the macro-average recall across classes. Empty cells and values associated with the New Datasets were excluded from the computation of placement-specific average metrics.

### External Validation Results

External validation confirmed high precision for Hand-held (94.2%), Lower-Back (100%), and Back/Front pockets (94.7%) but highlighted challenges with Shoulder Bag and Coat Pocket (Figure 5), achieving precision lower than 50%. Classification accuracy changed across different morbidities (0.71 - 0.85), walking speeds (0.79 - 0.87), activities (0.63 - 0.94), age groups (0.76 - 0.85) and genders (0.87 - 0.95), while neglectable effects were observed for the other factors (Figure 6).

**Figure 6.**
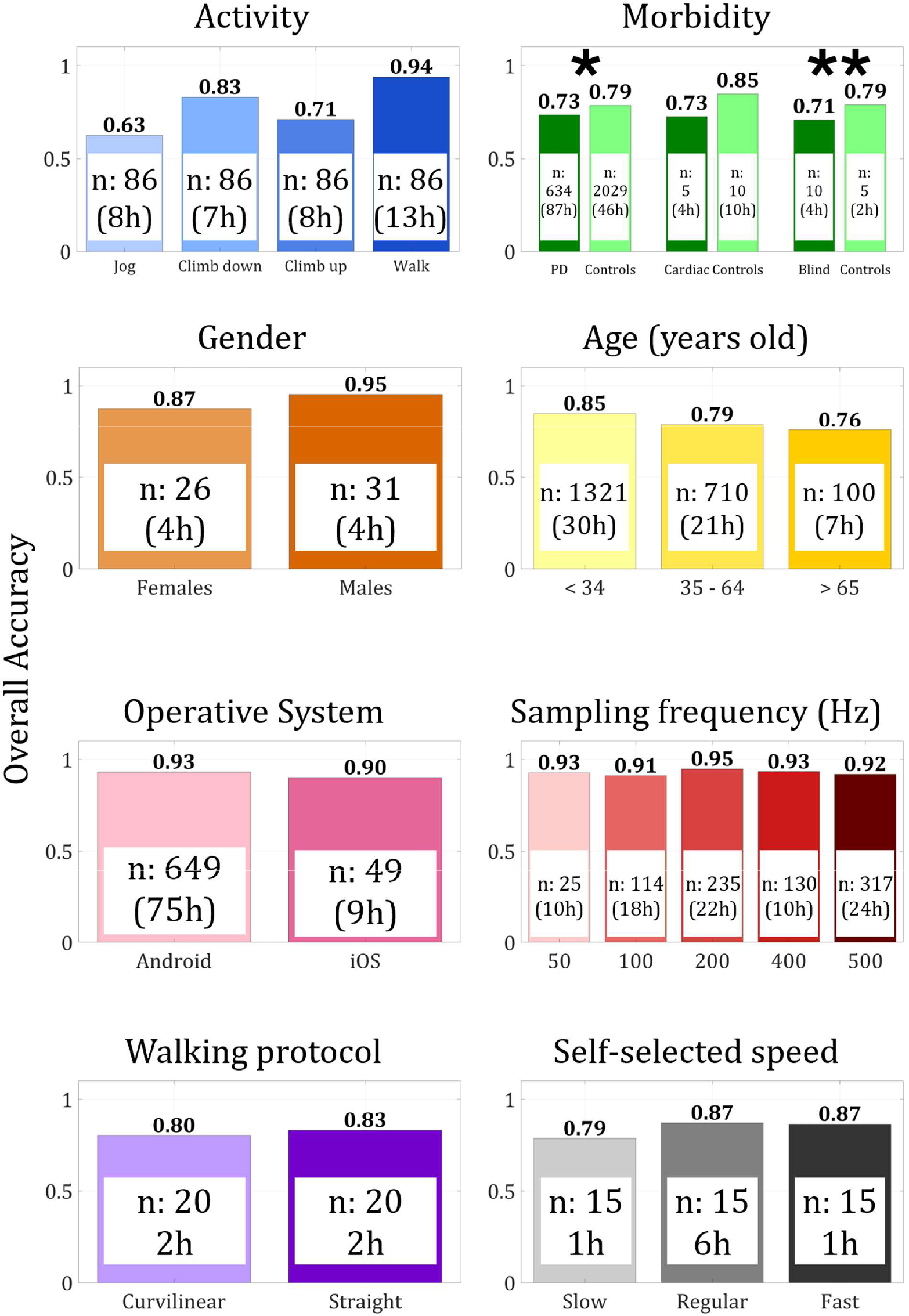
For each factor, accuracy of each group is reported on top of the corresponding bar. Dimension of the samples used in each group was quantified by the number of participants (n) and the hours of data (h), both reported within the corresponding bar. *Results for the effect of medication in Parkinson’s disease patients (mPower dataset) and **walking support type (cane or guide dog) in blind participants (WAW dataset) are reported in the Discussion.

## 4. Discussion

This study aimed to evaluate the robustness of smartphone placement recognition during real-world walking using heterogeneous datasets. The underlying hypothesis of the study was that inertial signals recorded by smartphones can be used to infer the device’s location. The study confirmed the feasibility of such task; however, it also highlighted the importance of heterogeneity of training data to achieve robustness against confounding factors such as morbidity, walking speed, concurrent activities, age, and gender.

Internal validation was conducted on free-living data from six on-body placements. In addition, the largest benchmarking of smartphone placement recognition to date was performed, involving over 3,000 healthy and pathological participants and more than 270 hours of heterogeneous data across devices, protocols, and gait characteristics. Finally, an extensive analysis of factors influencing recognition accuracy was carried out, providing practical guidance for experimental design and future digital mobility research.

Performance obtained by the model in controlled conditions (New Laboratory Dataset) was in line with similar machine-learning-based studies reporting overall accuracies in the range of 80–87% [31], [35], [36], [37]. Reduced performance observed in the 6-class setting was primarily driven by misclassification among pocket-related placements, attributable to the strong similarity of inertial signal characteristics recorded in these biomechanically comparable locations. This was confirmed by external validation, as aggregating pocket classes consistently improved performance: in the case of OxIOD, merging front and back placement classes led to 85% recall improvement.

Recognition precision and recall were highest for datasets including healthy participants in semi-controlled conditions (CHAD, MSR, OxIOD, RHAR, SAD, FDA, PAR, WDE), and decreased in free-living conditions (RuDaCop, mPower), although accuracy remained around 75% in the 4-class setting. For instance, RuDaCop, which captures unstructured free-living daily activities without supervision, and mPower, where participants performed the walking test independently at home following only app-based instructions, showed substantially lower recall for pocket classes. Accuracy of pocket class improved in Parkinson’s disease patients following medication (from 71.4% to 75.1%), suggesting that transient enhancements in gait regularity can positively influence classifier performance. Reduced accuracy was observed for WAW, which included blind individuals with markedly altered gait patterns and the use of walking aids or guide dogs. Using a guide dog was associated with lower recognition accuracy (68.7%) than using a walking cane (71.6%), likely because the altered arm and body movements required to handle the dog introduced additional variability in the smartphone’s motion patterns. In RIDI and RuDaCop lower performance was observed, due to the inclusion of shoulder-bag instances, which increased false positives. These findings suggest that bag placements, associated with greater variability and influenced by user habits, remain challenging for smartphone placement recognition classifiers.

The smartphone placement recognition classifier was trained with data collected during level walking, the most prevalent daily-life activity and the one of greatest relevance for mobility assessment. As part of the factor analysis, model robustness was further examined by evaluating how deviations from the training conditions—such as different locomotor activities and walking speeds—affected recognition performance (Figure 6): accuracy was high during walking (∼94%) but declined for stairs (71–83%) and, more markedly, for jogging (63%), in accordance with previous findings [29], [38]. The model performance was partially influenced by walking speed, with accuracy dropping from 0.87 (regular and faster speed) to 0.79 (slow speed), in line with previous studies that identified walking speed as a confounder in real-world digital mobility assessment [2]. This suggests that, at reduced speeds, gait features may differ less markedly between placements from those at higher speeds. Notably, the model was trained on real-world data likely containing fewer slow-walking strides, particularly compared with the deliberately simulated slow-speed trials in the Laboratory Custom dataset. Incorporating slow-walking data during training could therefore help mitigate this performance drop.

Morbidity affected smartphone placement recognition performance. Pathological cohorts generally exhibited slightly lower recognition accuracy compared to healthy controls. In the mPower dataset, the difference in accuracy between PD subjects (73%) and controls (79%) were likely attributable to the characteristics of Parkinsonian gait (rigidity, slowness, irregularity). These results must also be contextualised by the fully unsupervised 6-month ResearchKit protocol of the mPower study, which introduced ecological complexity and uncertainty regarding the actual device placement, as participants were not constrained to a specific pocket. Similarly, in the WAW dataset, blind subjects (71%) showed lower accuracy than sighted controls (79%). This dataset’s realistic protocol, involving aids, guide dogs, and environmental interactions (e.g., bumps, doors), likely contributed to this gap and to a general deterioration of performance with respect to other datasets. The largest cohort-wise difference was in the IPos dataset between cardiac patients (73%) and controls (85%), suggesting that substantial differences in gait characteristics influenced the signal features.

Small but noticeable differences in accuracy emerged across gender (female: 0.87, male: 0.95) and age groups (youth: 85%, adults: 79%, elderly: 76%). Although the model demonstrated overall robustness, generalizing reasonably well to women and older adults despite being trained predominantly on young, male participants, the slight performance deterioration aligns with the demographic composition of the training set.

Regarding hardware, no substantial accuracy differences emerged based on the device’s operating system (Android: 93%, iOS: 90%). Likewise, the original sampling frequency (which was resampled to 100 Hz pre-analysis) did not have a significant effect, suggesting the framework is robust to data collected at different frequencies and that resampling did not introduce significant signal variations in the calculated features. Finally, the adopted training strategy, which used free-living data inclusive of curvilinear motion, was effective for generalizing non-linear walking conditions, as no significant differences in accuracy were observed between straight (83%) and curvilinear (80%) walking.

The present findings show that smartphone placement can be reliably recognized in real-world walking conditions, but also indicate that performance is mainly driven by classification granularity and affected by contextual and subject-specific factors, with limited influence of hardware-related variability.

A primary weakness of this study was the difficulty in recognizing the Shoulder Bag placement, due to the high variability in bag type, phone placement, and carrying style across datasets. While internal validation benefited from the use of a standardised bag, performance deteriorated on heterogeneous external data. Future fine-tuning with more diverse samples may mitigate this issue, as well as strategies based on recognition of unknown classes [13]. The model also struggled to discriminate fine-grained pocket placements (Front, Back, Coat), reflecting their biomechanical similarity. As confirmed by results, aggregating them into a single Pocket class yielded substantially more reliable performance and is often sufficient for mobility-assessment applications where gait outcomes remain equivalent across pocket subtypes.

Across public datasets, the Lower-Back placement exhibited lower recall but consistently high precision. While reduced recall indicates occasional misclassification of Lower-Back segments, this is tolerable in clinical applications with standardized sensor placement [2], where precision is the key requirement to ensure that only correctly localized data are processed by downstream digital mobility algorithms.

The main analysis was restricted to walking data, excluding running and inclined walking. Prior works have proposed hybrid frameworks that jointly infer activity and placement under the assumption that transitions between them are relatively infrequent [38]; however, this may not hold in domestic contexts with frequent placement changes. A more robust solution for smartphone placement recognition during gait is to precede smartphone placement recognition with a gait sequence detection module, constraining classification to stable walking periods. This approach is supported by evidence showing that position-independent and position-aware activity recognition achieve comparable performance [35], indicating that activity classification can be applied effectively even without knowledge of phone placement. Moreover, while placement critically affects advanced mobility outcomes such as stride length [33], its impact on gait-sequence detection is limited, as demonstrated by studies where gait sequence detection algorithms developed for pelvic sensors were successfully applied to wrist-worn data [39].

Recognition from the Hand placement was evaluated only for natural arm swing, excluding real-world hand-use modes such as phoning, texting, scrolling, or holding bags. Prior work shows these modes can be classified, suggesting that expanding the taxonomy of hand motions would improve ecological validity and enable more flexible deployment [40].

Finally, training on the full 6-class taxonomy allowed post hoc evaluation at different levels of granularity without loss of information. However, training directly on aggregated classes could potentially improve performance by reducing ambiguity among biomechanically similar placements and simplifying decision boundaries. Training relied on data from a limited number of young, healthy participants. As shown in this work, age, gender, or pathology influence smartphone placement recognition accuracy, underscoring the need for more heterogeneous training sets to ensure generalization across populations.

Overall, these findings highlight the importance of developing smartphone placement recognition frameworks trained on datasets that reflect the breadth of real-world scenarios, including diverse speed profiles, pathological gait, and balanced demographic distributions.

## Conclusions

Accurate identification of smartphone placement is fundamental for reliable digital mobility assessment, as many gait measures depend on device placement. This study demonstrates that smartphone placement recognition can achieve robust performance across a wide range of real-world conditions, but also reveals key challenges related to uncontrolled environments, population heterogeneity, and fine-grained placement granularity, supporting the need for context-aware analysis approaches to ensure reliable deployment in real-world applications. This work aims to provide a roadmap for future smartphone placement recognition frameworks that are more generalizable and clinically usable.

## CRediT authorship contribution statement

**Paolo Tasca**: Writing – review & editing, Writing – original draft, Visualization, Validation, Software, Methodology, Investigation, Formal analysis, Data curation. **Giorgio Trentadue**: Writing – review & editing, Visualization, Validation, Software, Methodology, Investigation, Formal analysis, Data curation. **Ellen Buckley:** Writing – review & editing, Visualization, Supervision, Resources, Project administration, Methodology, Funding acquisition, Formal analysis, Conceptualization. **Shaoxiong Sun:** Writing – review & editing, Software, Visualization, Supervision, Methodology, Formal analysis, Conceptualization. **Michael Long:** Writing – review & editing, Supervision, Methodology, Formal analysis, Conceptualization. **Neil Ireson:** Writing – review & editing, Visualization, Supervision, Methodology, Formal analysis, Conceptualization. **Fabio Ciravegna:** Writing – review & editing, Software, Data curation, Methodology, Formal analysis, Conceptualization. **Vitaveska Lanfranchi:** Writing – review & editing, Visualization, Supervision, Resources, Project administration, Methodology, Funding acquisition, Formal analysis, Conceptualization. **Andrea Cereatti**: Writing – review & editing, Visualization, Supervision, Resources, Project administration, Methodology, Funding acquisition, Formal analysis, Conceptualization.

## Declaration of competing interest

The authors declare that they have no known competing financial interests or personal relationships that could have appeared to influence the work reported in this paper.

## Acknowledgements

This study was awarded the “SIAMOC Best Paper 2025” supported by Società Italiana di Analisi del Movimento in Clinica (SIAMOC) and Open Access Publishing Fund of SIAMOC, Italy

## Notes

https://github.com/H-MOVE-LAB/smartphone-placement-recognition

### Competing Interest Statement

The authors have declared no competing interest.

## References

[1] J. Albites-Sanabria et al., «Walking into aging: real-world mobility patterns and digital benchmarks from the InCHIANTI Study», Npj Aging, vol. 11, fasc. 1, p. 60, lug. 2025, doi: 10.1038/s41514-025-00245-w.

[2] C. Kirk et al., «Mobilise-D insights to estimate real-world walking speed in multiple conditions with a wearable device», Sci. Rep., vol. 14, fasc. 1, p. 1754, gen. 2024, doi: 10.1038/s41598-024-51766-5.

[3] J. Tian, L. Cong, e H. Qin, «Mixed-Pose Positioning in Smartphone-Based Pedestrian Dead Reckoning Using Hierarchical Clustering», IEEE Trans. Instrum. Meas., vol. 72, pp. 1–12, 2023, doi: 10.1109/TIM.2023.3326249.

[4] S. Tao, H. Zhang, L. Kong, Y. Sun, e J. Zhao, «Validation of gait analysis using smartphones: Reliability and validity», Digit. Health, vol. 10, p. 20552076241257054, set. 2024, doi: 10.1177/20552076241257054.

[5] B. M. Zeleke et al., «Mobile phone carrying locations and risk perception of men: A cross-sectional study», PLOS ONE, vol. 17, fasc. 6, p. e0269457, giu. 2022, doi: 10.1371/journal.pone.0269457.

[6] Q. Tian, Z. Salcic, K. I.-K. Wang, e Y. Pan, «A Multi-Mode Dead Reckoning System for Pedestrian Tracking Using Smartphones», IEEE Sens. J., vol. 16, fasc. 7, pp. 2079–2093, apr. 2016, doi: 10.1109/JSEN.2015.2510364.

[7] O. Asraf, F. Shama, e I. Klein, «PDRNet: A Deep-Learning Pedestrian Dead Reckoning Framework», IEEE Sens. J., vol. 22, fasc. 6, pp. 4932–4939, mar. 2022, doi: 10.1109/JSEN.2021.3066840.

[8] R. Yang e B. Wang, «PACP: A Position-Independent Activity Recognition Method Using Smartphone Sensors», Information, vol. 7, fasc. 4, Art. fasc. 4, dic. 2016, doi: 10.3390/info7040072.

[9] H. Yan, Q. Shan, e Y. Furukawa, «RIDI: Robust IMU Double Integration», in Computer Vision – ECCV 2018, vol. 11217, V. Ferrari, M. Hebert, C. Sminchisescu, e Y. Weiss, A c. di, in Lecture Notes in Computer Science, vol. 11217., Cham: Springer International Publishing, 2018, pp. 641–656. doi: 10.1007/978-3-030-01261-8_38.

[10] J.-S. Lee e S.-M. Huang, «An Experimental Heuristic Approach to Multi-Pose Pedestrian Dead Reckoning Without Using Magnetometers for Indoor Localization», IEEE Sens. J., vol. 19, fasc. 20, pp. 9532–9542, ott. 2019, doi: 10.1109/JSEN.2019.2926124.

[11] Wang, Ye, Luo, Haiyong, Men, Zhao, e Ou, «Pedestrian Walking Distance Estimation Based on Smartphone Mode Recognition», 2019, Consultato: 1 dicembre 2025. [Online]. Disponibile su: https://www.mdpi.com/2072-4292/11/9/1140

[12] Q. Du, Z. Wang, Y. Kuang, Y. Yao, Q. Cao, e Y. Yang, «P2Net: A Two-Stage Personalized Pedestrian Dead Reckoning Based on Neural Networks», IEEE Sens. J., vol. 25, fasc. 3, pp. 5757–5768, feb. 2025, doi: 10.1109/JSEN.2024.3515173.

[13] N. Daniel, F. Goldberg, e I. Klein, «Smartphone Location Recognition with Unknown Modes in Deep Feature Space», Sensors, vol. 21, fasc. 14, Art. fasc. 14, gen. 2021, doi: 10.3390/s21144807.

[14] I. Klein, «Smartphone Location Recognition: A Deep Learning-Based Approach», Sensors, vol. 20, fasc. 1, p. 214, dic. 2019, doi: 10.3390/s20010214.

[15] A. Cereatti et al., «ISB recommendations on the definition, estimation, and reporting of joint kinematics in human motion analysis applications using wearable inertial measurement technology», J. Biomech., vol. 173, p. 112225, ago. 2024, doi: 10.1016/j.jbiomech.2024.112225.

[16] F. Salis et al., «A multi-sensor wearable system for the assessment of diseased gait in real-world conditions», Front. Bioeng. Biotechnol., vol. 11, apr. 2023, doi: 10.3389/fbioe.2023.1143248.

[17] F. Salis, S. Bertuletti, T. Bonci, U. Della Croce, C. Mazzà, e A. Cereatti, «A method for gait events detection based on low spatial resolution pressure insoles data», J. Biomech., vol. 127, p. 110687, ott. 2021, doi: 10.1016/j.jbiomech.2021.110687.

[18] A. Bayev, I. Gartseev, I. Chistyakov, A. Nikulin, A. Derevyankin, e M. Pikhletsky, RuDaCoP: The Dataset for Smartphone-based Intellectual Pedestrian Navigation. 2019. doi: 10.48550/arXiv.1908.03609.

[19] B. M. Bot et al., «The mPower study, Parkinson disease mobile data collected using ResearchKit», Sci. Data, vol. 3, fasc. 1, p. 160011, mar. 2016, doi: 10.1038/sdata.2016.11.

[20] S. Caramaschi, C. M. Olsson, E. Orchard, J. Molloy, e D. Salvi, «Assessing the Effect of Data Quality on Distance Estimation in Smartphone-Based Outdoor 6MWT», Sensors, vol. 24, fasc. 8, p. 2632, apr. 2024, doi: 10.3390/s24082632.

[21] C. Chatzaki, M. Pediaditis, G. Vavoulas, e M. Tsiknakis, «Human Daily Activity and Fall Recognition Using a Smartphone’s Acceleration Sensor», lug. 2017, pp. 100–118. doi: 10.1007/978-3-319-62704-5_7.

[22] C. Chen, P. Zhao, C. X. Lu, W. Wang, A. Markham, e N. Trigoni, «OxIOD: The Dataset for Deep Inertial Odometry», 20 settembre 2018, arXiv: arXiv:1809.07491. doi: 10.48550/arXiv.1809.07491.

[23] G. H. Flores e R. Manduchi, «WeAllWalk: An Annotated Data Set of Inertial Sensor Time Series from Blind Walkers», in Proceedings of the 18th International ACM SIGACCESS Conference on Computers and Accessibility, in ASSETS ’16. New York, NY, USA: Association for Computing Machinery, ott. 2016, pp. 141–150. doi: 10.1145/2982142.2982179.

[24] M. Malekzadeh, R. G. Clegg, A. Cavallaro, e H. Haddadi, «Mobile Sensor Data Anonymization», in Proceedings of the International Conference on Internet of Things Design and Implementation, apr. 2019, pp. 49–58. doi: 10.1145/3302505.3310068.

[25] M. Matey-Sanz, S. Casteleyn, e C. Granell, «Dataset of inertial measurements of smartphones and smartwatches for human activity recognition», Data Brief, vol. 51, p. 109809, dic. 2023, doi: 10.1016/j.dib.2023.109809.

[26] M. Shoaib, S. Bosch, O. D. Incel, H. Scholten, e P. J. M. Havinga, «Complex Human Activity Recognition Using Smartphone and Wrist-Worn Motion Sensors», Sensors, vol. 16, fasc. 4, p. 426, apr. 2016, doi: 10.3390/s16040426.

[27] M. Shoaib, S. Bosch, O. D. Incel, H. Scholten, e P. J. M. Havinga, «Fusion of Smartphone Motion Sensors for Physical Activity Recognition», Sensors, vol. 14, fasc. 6, pp. 10146–10176, giu. 2014, doi: 10.3390/s140610146.

[28] M. Shoaib, H. Scholten, e P. J. M. Havinga, «Towards Physical Activity Recognition Using Smartphone Sensors», in 2013 IEEE 10th International Conference on Ubiquitous Intelligence and Computing and 2013 IEEE 10th International Conference on Autonomic and Trusted Computing, Italy: IEEE, dic. 2013, pp. 80–87. doi: 10.1109/UIC-ATC.2013.43.

[29] T. Sztyler e H. Stuckenschmidt, «On-body localization of wearable devices: An investigation of position-aware activity recognition», in 2016 IEEE International Conference on Pervasive Computing and Communications (PerCom), mar. 2016, pp. 1–9. doi: 10.1109/PERCOM.2016.7456521.

[30] U.S. Food and Drug Administration, «Open-Access Wearables Dataset to Evaluate Factors Impacting Accuracy of Smartphone Gait Metrics», 2024. Consultato: 27 ottobre 2025. [Online]. Disponibile su: https://www.synapse.org/Synapse:syn51664250/wiki/622541

[31] I. Klein, Y. Solaz, e G. Ohayon, «Smartphone Motion Mode Recognition», Proceedings, vol. 2, fasc. 3, Art. fasc. 3, 2017, doi: 10.3390/ecsa-4-04929.

[32] Y. Freund, «A more robust boosting algorithm», 13 maggio 2009, arXiv: arXiv:0905.2138. doi: 10.48550/arXiv.0905.2138.

[33] I. Klein, Y. Solaz, e G. Ohayon, «Pedestrian Dead Reckoning With Smartphone Mode Recognition», IEEE Sens. J., vol. 18, fasc. 18, pp. 7577–7584, set. 2018, doi: 10.1109/JSEN.2018.2861395.

[34] C. Ding e H. Peng, «Minimum redundancy feature selection from microarray gene expression data», J. Bioinform. Comput. Biol., vol. 3, fasc. 2, pp. 185–205, apr. 2005, doi: 10.1142/s0219720005001004.

[35] D. Coskun, O. D. Incel, e A. Ozgovde, «Phone position/placement detection using accelerometer: Impact on activity recognition», in 2015 IEEE Tenth International Conference on Intelligent Sensors, Sensor Networks and Information Processing (ISSNIP), apr. 2015, pp. 1–6. doi: 10.1109/ISSNIP.2015.7106915.

[36] K. Fujinami, «On-Body Smartphone Localization with an Accelerometer», Information, vol. 7, fasc. 2, p. 21, giu. 2016, doi: 10.3390/info7020021.

[37] O. D. Incel, «Analysis of Movement, Orientation and Rotation-Based Sensing for Phone Placement Recognition», Sensors, vol. 15, fasc. 10, pp. 25474–25506, ott. 2015, doi: 10.3390/s151025474.

[38] S. A. Antos, M. V. Albert, e K. P. Kording, «Hand, belt, pocket or bag: Practical activity tracking with mobile phones», J. Neurosci. Methods, vol. 231, pp. 22–30, lug. 2014, doi: 10.1016/j.jneumeth.2013.09.015.

[39] F. Kluge et al., «Real-World Gait Detection Using a Wrist-Worn Inertial Sensor: Validation Study», JMIR Form. Res., vol. 8, fasc. 1, p. e50035, mag. 2024, doi: 10.2196/50035.

[40] M. Susi, V. Renaudin, e G. Lachapelle, «Motion Mode Recognition and Step Detection Algorithms for Mobile Phone Users», Sensors, vol. 13, fasc. 2, pp. 1539–1562, feb. 2013, doi: 10.3390/s130201539.

