## Supplementary Materials for "Smartphone Placement Recognition during Walking: Performance Determinants and Real-World Generalizability"

### S1. Confusion Matrix

This section reports the confusion matrix obtained for the internal Test Set (New Free-living dataset). The matrix summarizes the classification performance across all placement classes, highlighting the distribution of true and misclassified instances, along with the percentages of correctly and mis-classified instances of each class (Figure S1).

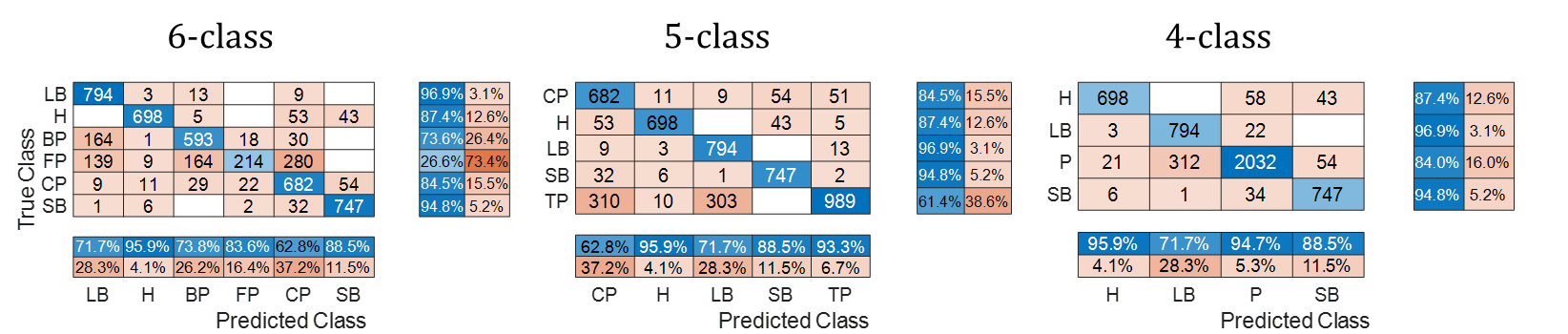

Figure S1 - Rows represent true classes and columns predicted classes. LB: Lower-Back; H: Hand-held; SB: Shoulder Bag; CP: Coat Pocket; BP: Back Pocket; FP: Front Pocket.

The confusion matrix illustrates the structure of misclassifications, showing that errors are predominantly concentrated among pocket-related placements, while Hand-held and Lower-Back placements are more distinctly recognized. As expected, the number of misclassified instances was reduced by merging the back pocket class to the front pocket class (5-class classification task), and subsequently to the coat pocket class (4-class classification task).
